# RNA-dependent RNA polymerase speed and fidelity are not the only determinants of the mechanism or efficiency of recombination

**DOI:** 10.1101/769224

**Authors:** Hyejeong Kim, Victor D. Ellis, Andrew Woodman, Yan Zhao, Jamie J. Arnold, Craig E. Cameron

## Abstract

Using the RNA-dependent RNA polymerase (RdRp) from poliovirus (PV) as our model system, we have shown that Lys-359 in motif-D functions as a general acid in the mechanism of nucleotidyl transfer. A K359H (KH) RdRp derivative is slow and faithful relative to wild-type enzyme. In the context of the virus, RdRp-coding sequence evolves, selecting for the following substitutions: I331F (IF, motif-C) and P356S (PS, motif-D). We have evaluated IF-KH, PS-KH, and IF-PS-KH viruses and enzymes. The speed and fidelity of each double mutant are equivalent. Each exhibits a unique recombination phenotype, with IF-KH being competent for copy-choice recombination and PS-KH being competent for forced-copy-choice recombination. Although the IF-PS-KH RdRp exhibits biochemical properties within twofold of wild type, the virus is impaired substantially for recombination in cells. We conclude that there are biochemical properties of the RdRp in addition to speed and fidelity that determine the mechanism and efficiency of recombination. The interwoven nature of speed, fidelity, the undefined property suggested here, and recombination makes it impossible to attribute a single property of the RdRp to fitness. However, the derivatives described here may permit elucidation of the importance of recombination on the fitness of the viral population in a background of constant polymerase speed and fidelity.

**Significance:** The availability of a “universal” method to create attenuated viruses for use as vaccine strains would permit a rapid response to outbreaks of newly emerging viruses. Targeting RdRp fidelity has emerged as such a universal approach. However, because polymerase fidelity and speed are inextricably linked, the effort to attribute the attenuated phenotype to a single biochemical property of the RdRp may be futile. Here, we show that this circumstance is even more complex. We provide evidence for the existence of a biochemical parameter that combines with fidelity and speed to govern the mechanism and/or efficiency of recombination. We conclude that the field will be served best by continued emphasis on discovery of manipulatable functions of the RdRp instead of debating the importance of individual properties.

## Introduction

In spite of the substantial resources that have been allocated by the National Institute of Allergy and Infectious Diseases and the Centers for Disease Control and Prevention to support prediction of emerging viral pathogens (https://www.niaid.nih.gov/research/emerging-infectious-diseases-pathogens), viral outbreaks over the past few decades have been caused by viruses for which surveillance was not considered a priority. Rapid response to an outbreak caused by an unexpected viral pathogen requires, minimally, the existence of broad-spectrum, antiviral therapeutics. Prevention requires the availability of vaccines, development of which could take years. Indeed, approved vaccines still do not exist to prevent infections by West Nile virus or severe acute respiratory syndrome (SARS) coronavirus, and these outbreaks occurred more than one decade ago.

Because all RNA viruses encode an RNA-dependent RNA polymerase (RdRp) with conserved function and mechanism, this enzyme has emerged as an attractive target for development of broad-spectrum therapeutics (1-3) and a target for function/mechanism-based strategies for viral attenuation (4-9). One function/mechanism of the RdRp that has been targeted most is that required for faithful incorporation of nucleotides (4-9). Changing RdRp fidelity decreases or increases the genetic variation of the viral population, which, in turn, decreases fitness and virulence of the viral population (10-15).

Known RdRp variants exhibiting a high-fidelity phenotype can also exhibit a reduced speed of nucleotide incorporation, at least at the biochemical level (16). Replication speed is also a determinant of viral fitness and virulence (17). So, is it the increased fidelity or decreased speed of nucleotide addition that gives rise to the attenuated phenotype? Further complicating the fidelity-versus-speed question are the recent observations that changes to fidelity also have consequence for the efficiency of recombination (18-23). Increased RdRp fidelity decreases recombination efficiency and vice versa (18-23). It will likely be impossible to attribute a single biochemical property of the RdRp to biological outcome.

The most extensively studied PV fidelity mutants encode RdRps with amino acid substitutions at sites remote from the active site. Constructing equivalent substitutions conferring equivalent phenotypes in RdRps other than PV is difficult if not impossible. Our laboratory has therefore pursued active-site-based strategies to manipulate the fidelity, speed, and/or recombination efficiency of the RdRp (24). We have shown that a lysine (Lys-359 in PV) present in conserved structural motif D of the RdRp contributes to the efficiency of nucleotidyl transfer (24). A PV mutant encoding a K359R RdRp is attenuated but elicits a protective immune response in mice that is at least as robust as the immune response elicited by the type 1 Sabin vaccine strain (6).

In this study, we characterize a second motif-D mutant of PV (K359H). Unlike the K359R RdRp-encoding virus (6), K359H PV is genetically unstable and acquires mutations encoding two second-site amino acid substitutions after a few passages in cell culture. Together, the two substitutions restore all biochemical properties of the derivative to near wild-type levels. Individually, however, we observe differences in the mechanism (copy-choice vs. forced-copy-choice) and efficiency of recombination by each derivative, although each derivative exhibits equivalent speed and fidelity. We conclude that biochemical properties in addition to speed and fidelity must exist and contribute to both the mechanism and efficiency of recombination. The desire to attribute single, biochemical properties of the viral RdRp to fitness, virulence, and pathogenesis may be futile.

## Results

### K359H PV requires two second-site substitutions to restore a “wild-type” growth phenotype

Most studies of RdRp fidelity have benefited from the selection of derivatives that were either more or less sensitive to a mutagenic nucleoside (25-30). Almost invariably, the derivatives changed residues remote from the catalytic site, thus using an allosteric mechanism to perturb fidelity. Among the most famous of these is the G64S substitution in PV RdRp (15, 16, 26). Many years ago, our laboratory showed that the active site of all polymerases contain a general acid, Lys-359 in the case of PV RdRp, that protonated the pyrophosphate leaving group during nucleotidyl transfer, thereby increasing the catalytic efficiency of RdRp (24). Substitutions of Lys-359 in PV RdRp give rise to changes in fidelity (6, 17). The arginine substitution of Lys-359 (K359R RdRp) and the histidine substitution (K359H RdRp) catalyze nucleotidyl transfer at a rate 10-fold lower than wild type (24). The impact of the arginine substitution of Lys-359 on PV fidelity and its potential application to vaccine development have been described (6).

Biological studies of K359H PV were not reported, because this virus was not genetically stable. When PV is rescued from in vitro transcribed RNA, four passages are required for the genetic diversity of the viral population to come to equilibrium (10). During this time for K359H PV, mutations were observed in RdRp-coding sequence that changed the plaque phenotype of the virus. One change was in motif C, I331F, and the other change was in motif D, P356S (**Figs. 1A** and **1B**). In order to compare the impact of these substitutions on PV genome replication without complication of reversion, we engineered the various substitutions into a subgenomic replicon, producing luciferase as an indirect measure of viral RNA produced. Experiments in the presence of guanidine hydrochloride (GuHCl) report on translation of transfected RNA in the absence of replication (**Fig. 1C**). Relative to WT PV, K359H was the most debilitated, with replication failing to reach an end point at 10 h post-transfection (**Fig. 1C**). Each double mutant was markedly better than K359H alone, but the addition of I331F conferred a greater replication advantage than P356S (**Fig. 1C**). The triple mutant replicated even faster but still exhibited a significant reduction in the rate of replication relative to WT (**Fig. 1C**).

**Figure 1:**
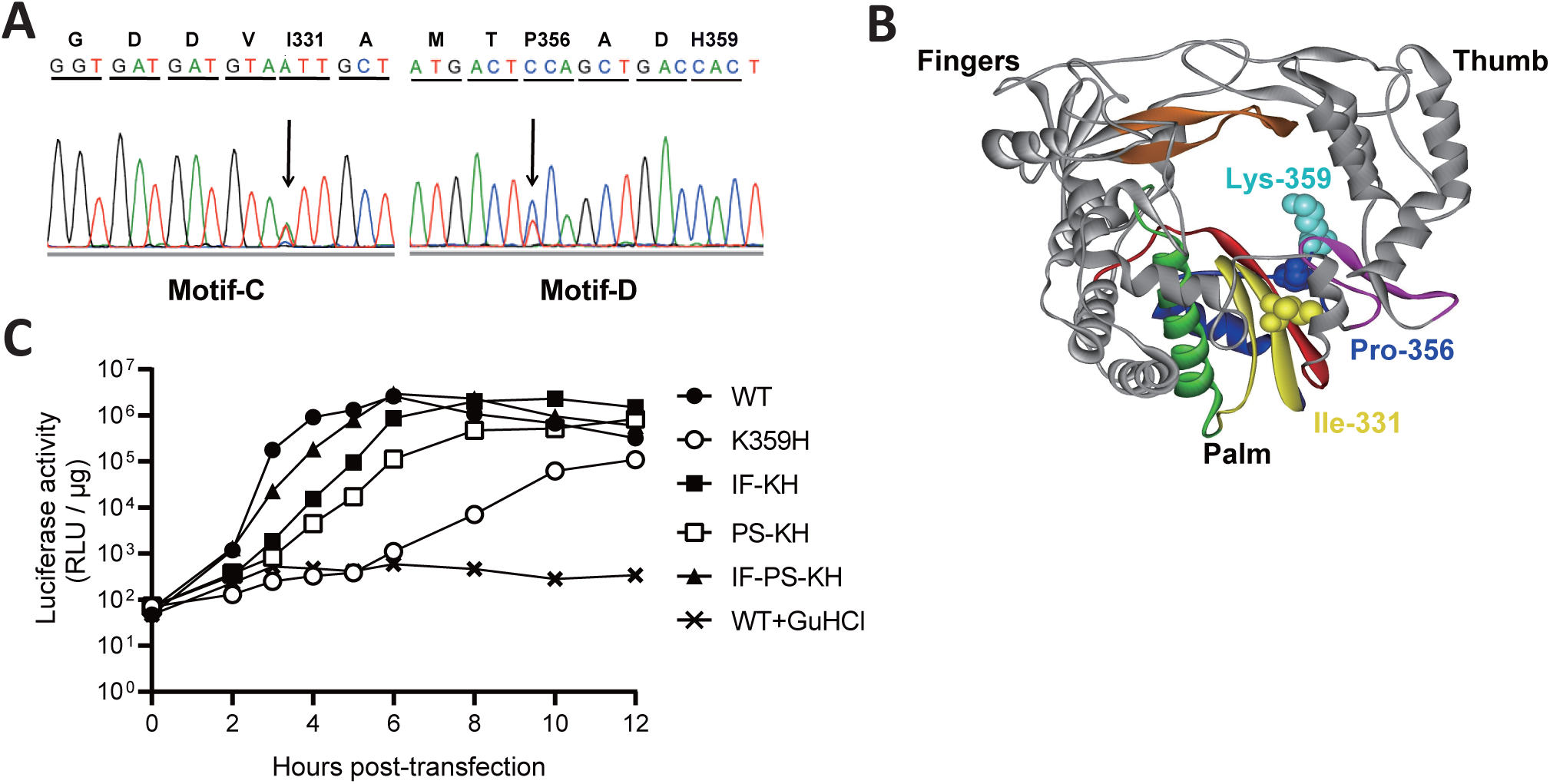
Serial passage of K359H PV in cell culture leads to additional changes to RdRp-coding sequence. (**A**) Changes in RdRp-coding sequence after serial passage of K359H PV. RNA was isolated from passage 2, converted to cDNA, and amplified by PCR. The electropherogram of the sequenced PCR product is shown, revealing two amino acid changes: I331F (motif C) and P356S (motif D). (**B**) Location of I331, P356, and K359 in structure of PV RdRp. Palm, fingers, and thumb subdomains are shown. Conserved structural motifs are colored: A, *red*; B, *green*; C, *yellow*; D, *blue*; E, *purple*; F, *orange*; G, *black*. (**C**) Replication phenotypes observed for K359H PV and KH-containing PV mutants by using a replicon assay. Replication was monitored using a subgenomic replicon firefly luciferase. Luciferase specific activity is reported in relative light units (*RLU*) per microgram of total protein in the extract as a function of time post-transfection. Shown is one representative data set.

### Characterization of the double- and triple-mutant viruses and their polymerases

At multiplicities of infection of one or higher, the double- and triple-mutant viruses were stable for multiple passages, thus permitting us to characterize the biological properties of these viruses. We have adopted the following nomenclature to refer to the various mutant PVs and their corresponding RdRps: I331F-K359H, IF-KH; P356S-K359H, PS-KH; and I331F-P356S-K359H, IF-PS-KH. Interestingly, the growth properties of the viruses were not as expected based on the experiments with the replicon. Each mutant virus exhibited the same delay relative to WT virus prior to the first detection of infectious virus after four hours post-infection (hpi) (**Fig. 2A**). Thereafter, growth of each virus was far more distinct than observed for the replicon, with PS-KH PV much slower than IF-KH PV and IF-PS-KH PV was substantially faster than both double mutants (**Fig. 2A**). The differences in outcome could reflect the differences in replication efficiency and/or a direct consequence of the substitutions in 3D-coding sequence on virus assembly or spread caused by changes to 3CD protein or 3D-containing precursor protein (31, 32).

**Figure 2:**
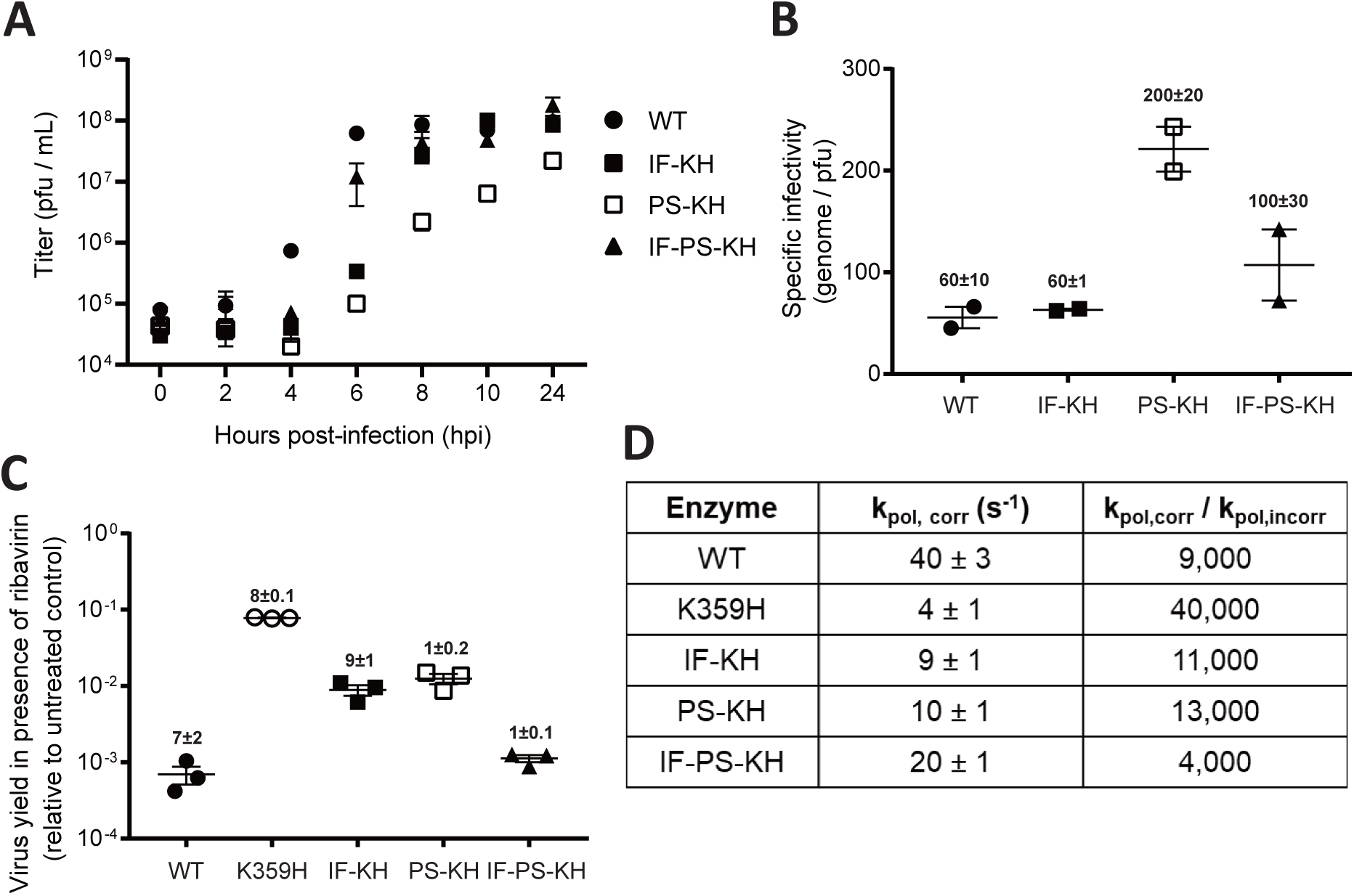
Biological and biochemical characterization of variants of K359H PV and the RdRp reveal that both substitutions are required to achieve a near wild-type phenotype. **(A**) One-step growth analysis. Cells were infected at MOI 10 with the following PVs: WT, IF-KH, PS-KH, and IF-PS-KH. Viral titer (pfu/mL) was plotted as a function of time post-infection. Duplicate infected samples were used for plaque assays. Error bars indicate SEM (*n = 2*). **(B)** Specific infectivity. Virus was isolated 24 h post-infection and used for qRT-PCR to determine genomes/mL or plaque assay to determine pfu/mL, with the quotient yielding specific infectivity, genomes/pfu. **(C)** Ribavirin sensitivity. HeLa cells were infected at a MOI 0.1 with each PV in the presence of 600 µM ribavirin. After a 24-h incubation at 37 °C, virus was isolated and used for plaque assay. Indicated is the titer of virus recovered in the presence of ribavirin normalized to that recovered in the absence of ribavirin. Solid bar indicates the mean of each virus yield. Error bars indicate SEM (*n = 3*). (**D**) Biochemical analysis. The rate constant for incorporation of a single correct nucleotide, ATP, (*k*_pol,corr_)) and a single incorrect nucleotide, GTP, (*k*_pol,incorr_) by each PV RdRp was performed as previously described (17). Data are reported using one significant figure. The error reported is the standard error from the fit of the data to a single exponential (17).

Our studies of RdRp fidelity mutants in cell culture have highlighted the fact that the use of the plaque-forming unit (PFU) as a measure of virus concentration will mask changes in the specific infectivity of the viral RNA (10). As a result, our experiments generally use genomes instead of PFU as the unit of measure (10). In doing so, we can use genomes/PFU as a measure of the specific infectivity of the virus and surrogate for virus fitness. The instability of KH PV precluded rigorous, quantitative analysis of this virus, but at least 10-fold more KH PV than WT PV was required to observe a comparable number of plaques. Both substitutions were able to increase the efficiency of plaque formation relative to KH PV (**Fig. 2B**). Indeed, the specific infectivity of IF-KH PV was equivalent to WT. The addition of PS to IF-KH reduced the efficiency of plaque formation; the specific infectivity of IF-PS-KH PV was reduced by twofold relative to IF-KH PV (**Fig. 2B**). Therefore, the exaggerated behavior of PS-KH PV in the PFU-based growth assay relative to the replicon assay likely reflects an additional defect to virus assembly and/or spread.

A change to the specific infectivity of the viral RNA caused by increased mutational load—that is, reduced fidelity of the PS-KH RdRp, would also explain the observed reduction in the specific infectivity of PS-KH PV relative to the other mutant PVs. To test this possibility, we evaluated the sensitivity of each virus population to growth in the presence of ribavirin. We reasoned that the mutagenic activity of ribavirin would exhibit the greatest negative impact on viruses with a mutator phenotype as described previously (10). Relative to WT, K359H PV exhibited the highest fidelity based on the 2-log difference in sensitivity to ribavirin (**Fig. 2C**). IF-KH and PS-KH PVs also exhibited a higher fidelity than WT PV (**Fig. 2C**). Importantly, both mutants exhibited essentially equivalent fidelity phenotypes (**Fig. 2C**), consistent with the suggestion above that the reduced efficiency of plaque formation observed for PS-KH PV is likely related to impairment of virus assembly and/or spread. Finally, evaluation of IF-PS-KH PV revealed equivalent sensitivity of this mutant to ribavirin as observed for WT PV (**Fig. 2C**), suggesting that fidelity had been returned to normal.

We have previously reported biochemical properties of the RdRps for the repertoire of mutant viruses described above (17). Consistent with these reports, each substitution increases the speed and reduces the fidelity to create a biochemical phenotype on par with that observed for WT (**Fig. 2D**).

### Biochemical properties of the RdRp other than speed and fidelity contribute to the efficiency of recombination in cell culture

Based on the myriad RdRp fidelity mutants that have been reported to date, there appears to be a direct correlation between the rate of nucleotide addition (speed) and the fidelity of nucleotide addition (6, 10, 17, 22, 33). This correlation also extends to recombination efficiency (22). If this is the case, then the ongoing debate of speed versus fidelity as the key determinant of viral fitness will become even more complicated to resolve (17).

The current state of the art for evaluation of recombination in cell culture is based on the co-transfection of two viral (sub)genomic RNAs incapable of producing infectious virus (34). The donor RNA is a replication-competent, subgenomic RNA that encodes a luciferase reporter instead of the viral capsid (**Fig. 3A**). The acceptor RNA is a replication-incompetent, genomic RNA that has a defective cis-acting replication element, termed oriI (**Fig. 3A**) (34). Initiation of replication on the donor followed by a switch to the acceptor at a site after the oriI locus will yield an infectious genome that can be scored by plaque assay (**Fig. 3A**) (34).

**Figure 3:**
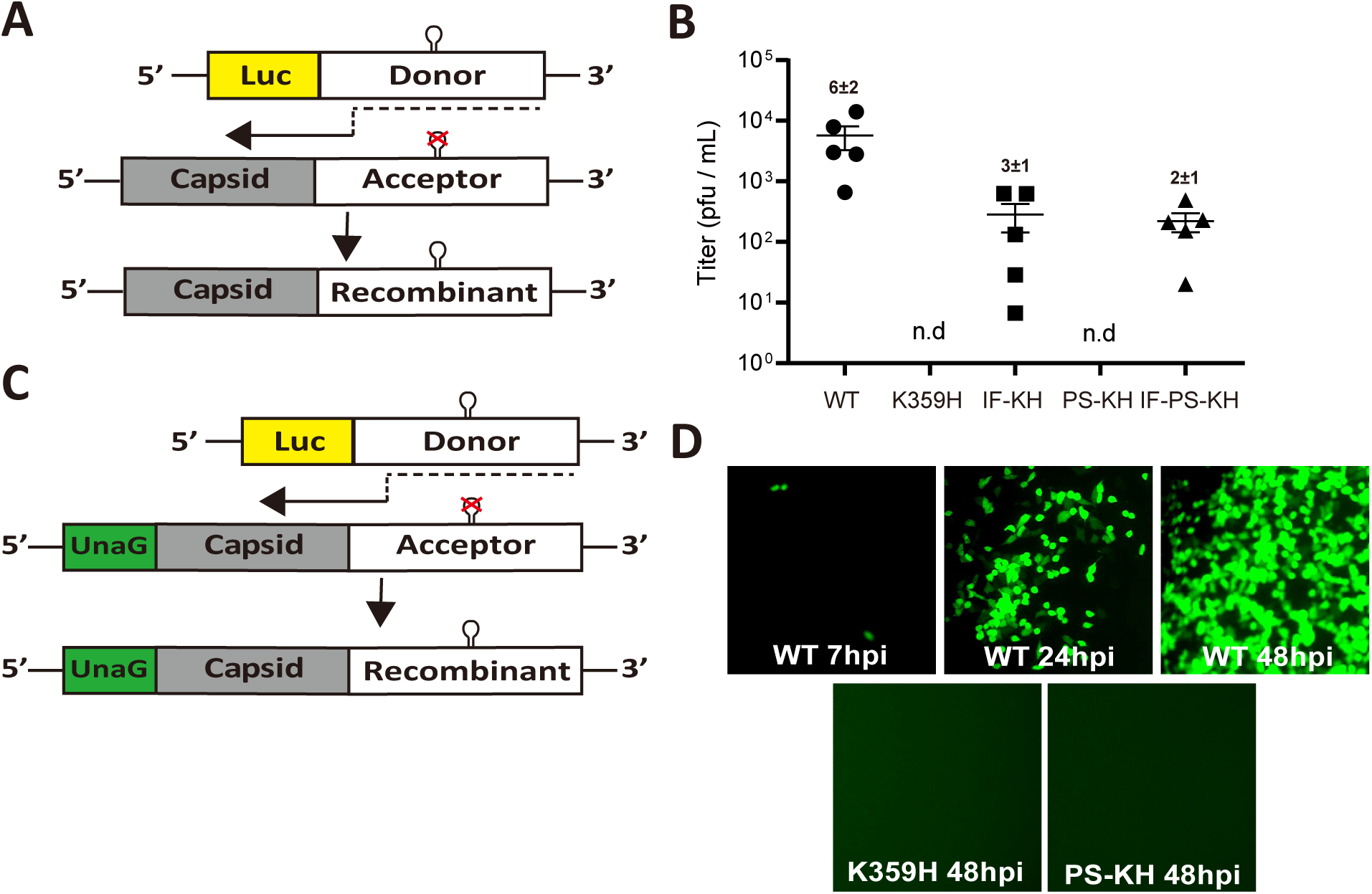
Parameters in addition to RdRp speed and fidelity contribute to recombination efficiency. **(A**) Schematic of the PV recombination assay used (34). Two RNAs are used: a replication-competent subgenomic RNA lacking capsid-coding sequence (donor RNA); and a replication-incompetent full-length genomic RNA with a defective cis-acting replication element (CRE, indicated by hairpin with defective version indicated by X) in 2C-coding sequence (acceptor RNA). Co-transfection of these RNAs produces infectious virus if recombination occurs. **(B**) The indicated PV RdRp was engineered into both donor and accepter RNA and co-transfected into a L929 mouse fibroblast cell line. Infectious virus produced by recombination in L929 cells was determined by plaque assay using HeLa cells. Each point shown is an independent experiment reflecting the average of three replicates. When plaques could not be detected, n.d. is indicated. Mean and SEM (n=5) are indicated. **(C**) Schematic of a modified PV recombination assay using an acceptor RNA expressing the green fluorescent protein, UnaG (35). Recombinants are scored by expression of green fluorescence instead of plaques, thereby increasing the sensitivity. **(D**) Infectious virus produced by recombination in L929 was scored for infectious virus in HeLa cells 7, 24, and 48 h post-infection for WT and at 48 h only for K359H and PS-KH.

We evaluated our panel of KH-containing PV mutants using this assay, and the outcomes were, in most cases, quite unexpected (**Fig. 3B**). KH PV was unable to produce viable recombinants, as expected for a high-fidelity RdRp (22). The first surprise was that IF-KH and PS-KH PVs did not exhibit the same phenotypes (**Fig. 3B**). While IF-KH PV produced viable recombinant virus, PS-KH PV did not (**Fig. 3B**). Based on the experiments performed above, these viruses and their polymerases replicate with comparable efficiency and fidelity (**Figs. 1A, 2C**, and **2D**). Even more surprising, however, was the observation that IF-PS-KH PV was impaired more than one log relative to WT PV in its ability to produce viable recombinants (**Fig. 3B**). The replication efficiency and fidelity of this virus and its polymerase were always within twofold of that observed for WT (**Figs. 1A, 2C**, and **2D**). Together, these results suggest that there are properties of the PV RdRp important for recombination that are not revealed by our existing biological and biochemical assays.

The one caveat of the recombination assay is that the recombinant viruses produced must be able to form plaques, which may mean that the recombinant virus must spread by a lytic mechanism. It was possible that PS-KH PV was defective for virus assembly and/or spread, at least lytic spread (**Fig. 2B**). In order to score for recombinant viruses that spread by either a lytic or non-lytic mechanism, we engineered the acceptor template to encode the UnaG (35) green fluorescent protein (**Fig. 3C**). The polyprotein was designed such that UnaG protein is released from capsid precursor protein by 3C protease activity. Observation of green cells were visible from a donor-acceptor pair producing WT RdRp as early as 7 hpi, with increases continuing over a 48-h period (**Fig. 3D**). With the sensitivity of this assay and the absence of a requirement for lytic spread, it is clear that PS-KH PV exhibits a recombination defect (**Fig. 3D**).

### Poly(rU) polymerase activity as a predictor of the efficiency of copy-choice recombination in cell culture

Twenty years ago, we showed that template switching was the primary mechanism of product formation when oligo(dT) or oligo(rU) were used to prime poly(rU) RNA synthesis on oligo(rA) or poly(rA) RNA templates (36). Briefly, polymerase engages a primed template, elongates that primer, disengages from the first template at some point during the elongation process to engage a second (acceptor) template and continue RNA synthesis (**Fig. 4A**) (36). This process occurs reiteratively, yielding products that are much greater than the average length of template used in the reaction (36).

**Figure 4:**
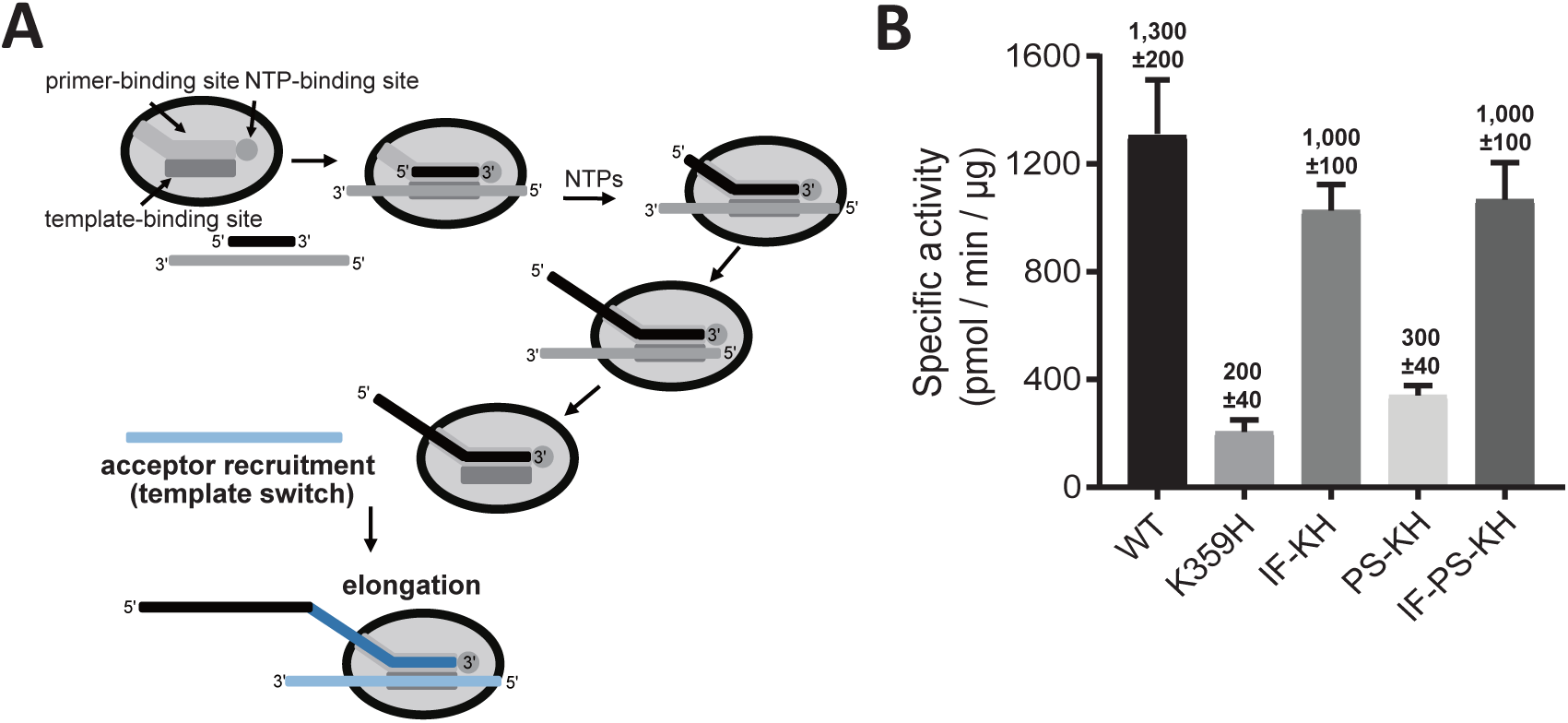
RdRp derivatives incapable of recombination in cells exhibit defects in copy-choice recombination (template switching) in vitro. **(A)** Products of reactions using oligo/poly(rA) as template reflect copy-choice recombination, which requires the template-switching activity of the RdRp (36). RdRp engages primed template and initiates RNA synthesis. The RdRp-nascent RNA complex moves to new template from internal positions or as a consequence of reaching the end of template, thus creating greater-than-template-length products. **(B)** Specific poly(rU) polymerase activity (pmol/min/µg) of each RdRp. WT and the indicated PV RdRp variants assessed by using a poly(rU) polymerase activity assay. Error bars represent SEM (*n = 3*).

The observation that only a subset of the KH mutants were competent for recombination, presumably by a copy-choice mechanism (37), provided the opportunity to determine the extent to which the poly(rU) polymerase activity can predict biological phenotypes. The congruence between the two experiments was remarkable (**Fig. 4B**). Both KH and PS-KH RdRps were impaired for poly(rU) polymerase activity, both IF-KH and IF-PS-KH RdRps exhibited near-WT levels of poly(rU) polymerase activity (**Fig. 4B**). These results further confirm template switching as the primary mechanism of poly(rU) polymerase activity and validate this assay as a screen for identification of RdRps with deficits in template switching.

### Polymerase determinants supporting efficient copy-choice recombination do not overlap completely with determinants supporting efficient forced-copy-choice recombination

Recently, our laboratory developed an assay for forced-copy-choice recombination (22, 38), inspired by an analogous assay developed to study template switching by the reverse transcriptase from human immunodeficiency virus (39-41). The concept of the assay and templates used are presented in **Fig. 5A**. The experimental design is diagrammed in **Fig. 5B**. Polymerase assembles on the primed template (sym/sub-U). In the presence of only the first nucleotide (ATP), the assembled complexes can be identified as the one-nucleotide-extended product (n+1). Addition of the remaining nucleotides in the absence or presence of the acceptor template will yield a strong-stop RNA product. In the presence of a complementary acceptor RNA, the strong-strop RNA product (donor) will be extended, creating a transfer product.

**Figure 5:**
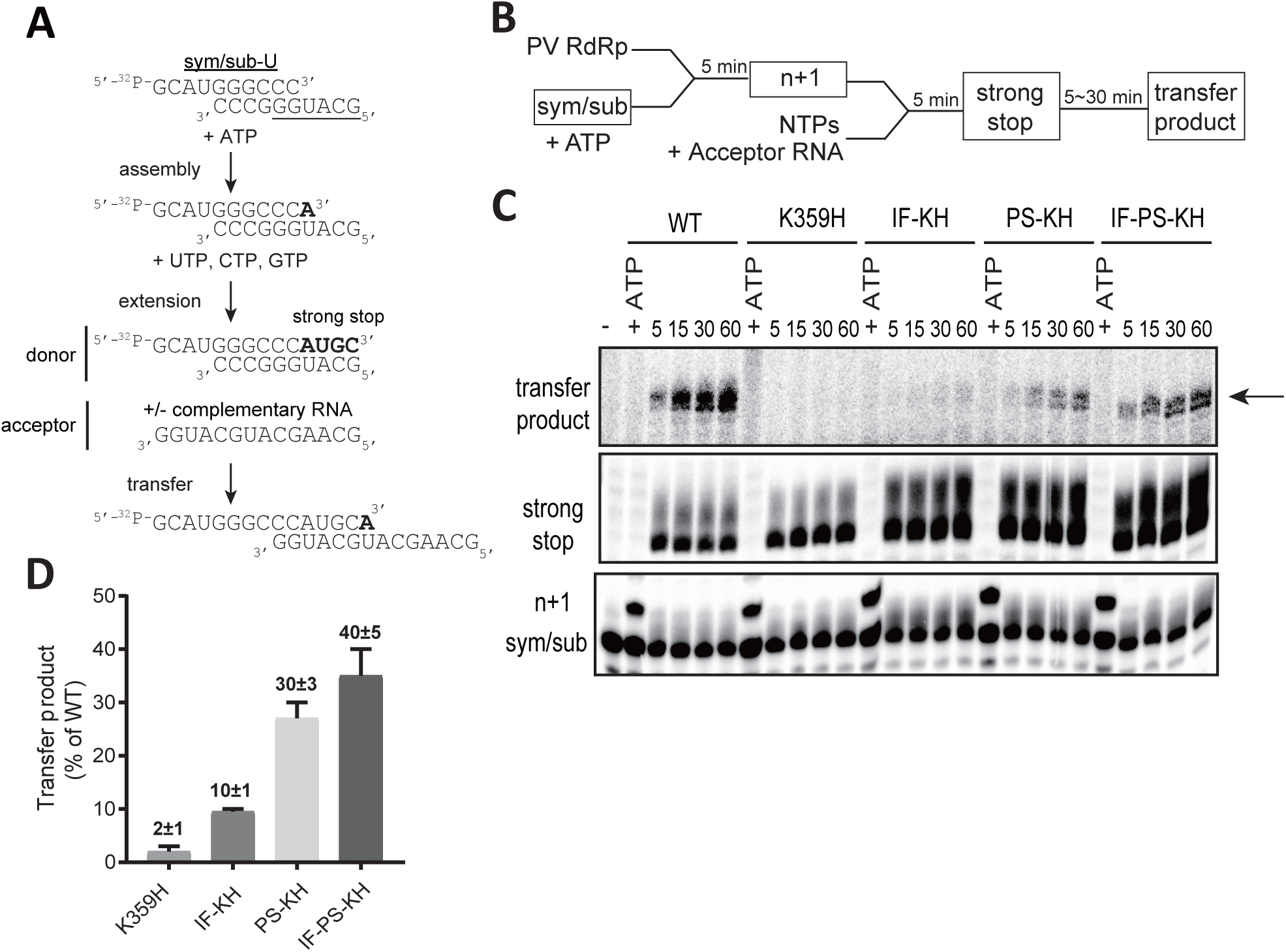
Properties of the RdRp required for copy-choice recombination in vitro are separable from those required for forced-copy-choice recombination in vitro. **(A)** The heteropolymeric, symmetrical, primed-template substrate (sym/sub) has been used to establish an assay for forced-copy-choice recombination in vitro (22, 38). RdRp assembles on sym/sub. ATP is added and incorporated to yield a stable elongation complex. It is this elongation complex that is monitored for extension and transfer. Transfer is strictly dependent on the presence of an acceptor RNA with complementarity to the 3’-end of donor RNA. **(B)** Schematic of the experimental design is indicated. Products in boxes are those observed and monitored by denaturing polyacrylamide gel electrophoresis. (**C**) Reaction products were resolved by electrophoresis and detected by phosphorimaging. The only regions of the gel with bands are shown; these correspond to the sym/sub primer, one-nucleotide-extended primer (n+1), four-nucleotides-extended product (strong stop) and non-templated addition of nucleotides to that product, and the transfer product. **(D)** Transfer products were quantified and are expressed as a percentage relative to the value observed for WT (The concentrations (µM) of n+1 formed for WT, K359H, IF-KH, PS-KH and IF-PS-KH are 0.22, 0.22, 0.32, 0.36 and 0.37, respectively. The concentrations (µM) of transfer product for WT, K359H, IF-KH, PS-KH and IF-PS-KH are 0.011, 0.00011, 0.0016, 0.0072 and 0.01, respectively. Overall, transfer efficiency ranges from 1% to 5%. Error bars represent SEM (n = 2).

We evaluated each KH-containing RdRp derivative in this assay. Substrate and product analysis are presented in **Fig. 5C**, with the quantitation of the transfer product relative to WT presented in **Fig. 5D**. KH RdRp was defective in this assay as well, as transfer product was not detected (**Fig. 5C**). Importantly, the failure to transfer was not a reflection of the inability to assemble or produce strong-stop donor RNA (**Fig. 5C**). In this assay, PS-KH RdRp outperformed IF-KH RdRp, and PS-KH and IF-PS-KH RdRp were essentially identical in activity.

We conclude that copy-choice and forced-copy-choice recombination use distinct mechanisms, requiring unique biochemical properties of the RdRp.

## Discussion

For more than a decade now, our laboratory and others have published studies asserting connections between viral RdRp fidelity and viral fitness (10-15). However, in the few instances in which RdRp derivatives exhibiting perturbed fidelity have been characterized, these derivatives also exhibit changes to the rate of nucleotide addition (6, 10, 17, 22, 33). Relative to wild-type polymerase, a higher fidelity polymerase is a slower enzyme, and a lower fidelity polymerase is a faster enzyme (6, 10, 17, 22, 33). Polymerase speed will clearly determine replication kinetics and therefore can contribute to viral fitness, as has been suggested recently (17). The most extensively characterized fidelity mutants of PV harbor an RdRp with a substitution located at a remote site that likely causes substantial collateral damage, further confounding the fidelity-versus-speed debate (4, 16). This study was motivated by the need to understand better the relationship between the biochemical properties of the viral RdRp and viral fitness, virulence, and pathogenesis. We have had in hand for a long time a PV mutant whose speed and fidelity were perturbed by changing an RdRp active-site residue, K359H (24). We did not publish this PV mutant until now because of its genetic instability (**Fig. 1A**). We realized that this genetic instability might actually represent an opportunity, as the biochemical phenotypes reverted by the second-site suppressors might highlight the biochemical properties driving viral fitness.

Lys-359 protonates the pyrophosphate leaving group during nucleotidyl transfer (24, 42). The K359H substitution will change the efficiency of that protonation event for two reasons. First, the pKa value of histidine is substantially lower than lysine and will therefore not be protonated to the same extent. Second, the distance between the nucleotide phosphates and the imidazole ring of the histidine will be greater than that of the amino group of lysine. It is likely that the distance is more of an issue, because K359R PV is stable and K359R RdRp exhibits the same reduction in catalytic efficiency as K359H RdRp (6). The changes of I331F and P356S in motifs C and D, respectively, may serve to move His-359 closer to the nucleotide phosphates.

Each substitution (IF and PS) individually increased the replication efficiency of K359H PV, with the combination of the two substitutions producing an additive outcome (**Fig. 1C**). Each substitution also decreased the fidelity of nucleotide addition in cells, as assessed by ribavirin sensitivity, to the same extent; the combination mutant yielded a virus with properties on par with wild type (**Fig. 2C**). The observed changes to replication speed and fidelity for the mutant viruses were explained by the biochemical properties of the corresponding RdRps (**Fig. 2D**). As observed in the past, both speed and fidelity appear to be correlated.

The substitutions at the active site of the polymerase do not only affect replication efficiency and population diversity but also impact other stages of the lifecycle. The kinetics of virus production for the different mutants did not correspond directly to the kinetics of replication (compare **Fig. 2A** to **Fig 1C**). Virus production was significantly delayed for PS-KH PV (**Fig. 2A**). This issue with virus production manifests as a substantial decrease in the specific infectivity of this virus (**Fig. 2B**). One possible explanation for this outcome is that the RdRp is encoded by the 3D region of the viral genome and this region is also a component of the PV 3CD protein. The 3CD protein has well established roles in aspects of the lifecycle before, during, and after replication, including virion morphogenesis (31, 32, 43, 44). Observations such as these highlight the additional level of complexity associated with establishing a cause-and-effect relationship between biochemical properties of the RdRp and fitness, virulence, and/or pathogenesis.

We and others have observed correlations between RdRp fidelity and recombination efficiency, with higher fidelity suppressing recombination and vice versa (18-23). An unexpected outcome of this study was the observation that IF-KH and PS-KH PVs exhibit substantially different propensities for recombination in cells (**Fig. 3B**), in spite of very similar speed and fidelity phenotypes in cells (**Fig. 1C** and **Fig. 2C**) and in vitro (**Fig. 2D**). PS-KH PV appeared to be completely incapable of supporting recombination in cells (**Fig. 3B**). Because the cell-based recombination assay requires the ability of virus to plaque, it was conceivable that the defect of PS-KH PV was more a reflection of spread than impaired recombination. We established a more sensitive fluorescence-based assay (**Fig. 3C**), which also indicated impaired recombination (**Fig. 3D**). The impact of the PS substitution on recombination was also quite evident in the context of IF-PS-KH PV. The biological and biochemical properties of this triple mutant are within twofold of wild type (**Figs. 1** and **2**), but the recombination efficiency of this mutant is down by 30-fold relative to WT PV (**Fig. 3B**). Together, these observations support the existence of biochemical properties of the RdRp other than speed and fidelity that are essential for recombination and impaired by changing Pro-356 to Ser. Further characterization of these mutants will be required to identify this undefined biochemical property.

Our laboratory has had a longstanding interest in the mechanism of recombination (18, 19, 22, 36, 38). We have shown that the high poly(rU) polymerase activity of PV RdRp derives from template switching during elongation, thus mimicking copy-choice recombination (**Fig. 4A**) (36). A second assay that we developed is based on template switching from the end of template, thus mimicking forced-copy-choice recombination (**Fig. 5A**) (22, 38). Based on the mutants and corresponding RdRp derivatives reported here, high fidelity impairs both types of recombination (see K359H in **Figs. 4B** and **5D**). Interestingly, unique RdRp determinants exist for each mode of recombination. PS-KH RdRp is impaired for copy-choice recombination (**Fig. 4B**); IF-KH is impaired for forced-copy-choice recombination (**Fig. 5D**). Impairment of recombination by PS-KH PV in cells is consistent with template switching as the primary mechanism of copy-choice recombination in cells, as demonstrated by Kirkegaard and Baltimore (37). The observation that IF-KH PV fails to exhibit a wild-type recombination phenotype (**Fig. 3B**) may suggest that forced-copy-choice recombination contributes to recombination in cells as well.

The current narrative of the characterized PV mutants impaired for recombination is that recombination acts as a mechanism to purge deleterious mutations in the viral population (45-47). Our study reveals several caveats for the interpretation of these previous studies. There are known and unknown biochemical properties of the RdRp that determine recombination efficiency, and there are at least two distinct mechanisms of recombination. Without understanding these complexities for the mutants under investigation, it is difficult to compare one study to another and may explain disparate observations made between laboratories (48, 49). Further analysis of the panel of mutants reported here may help to resolve the controversy but should illuminate how changes to the mechanism and efficiency of recombination impact viral evolution in a background of constant speed and fidelity.

Viral fitness, virulence, and pathogenesis are determined collectively by the structure, dynamics, and activity of all virus-encoded functions. The conserved structure, dynamics, and mechanism of the viral RdRp makes this enzyme an attractive target for development of attenuated viruses by changing conserved residues capable of perturbing conserved, biochemical function (6, 8, 10). As a field, we have focused on RdRp fidelity (6) but have come to the realization that RdRp speed should not be ignored (17). This study shows that speed, fidelity, and undefined biochemical properties of the RdRp exist and contribute to both the mechanism and efficiency of recombination (**Fig. 6**). This inextricable connection of the myriad biochemical properties of the RdRp precludes attribution of a single biochemical property to viral fitness, virulence, and pathogenesis (**Fig. 6**). The field will be served best by continued emphasis on discovery of manipulatable functions of the RdRp, and other viral enzymes, instead of debating the importance of individual properties.

**Figure 6:**
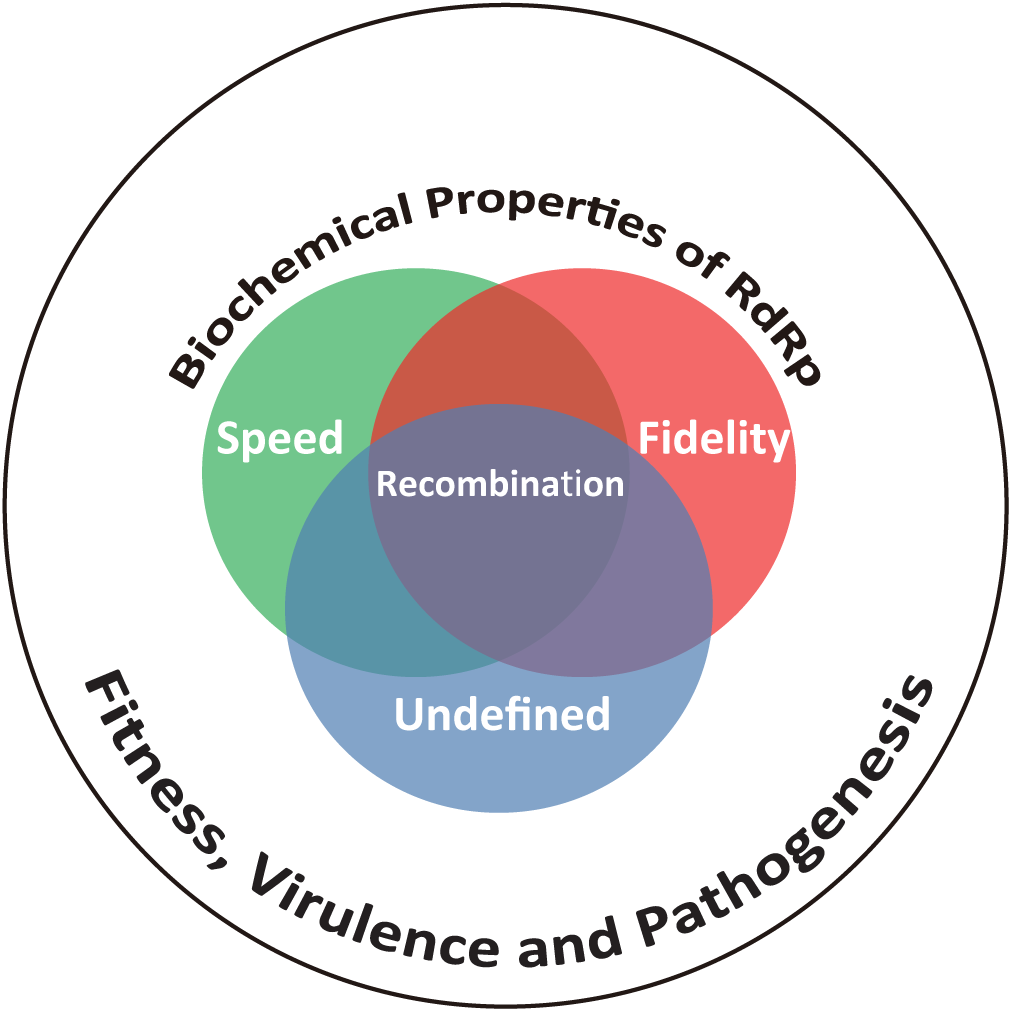
The biochemical properties of the RdRp are inextricably linked and collectively determine biological outcomes. PV RdRp is among the most extensively studied enzyme both in cells and in test tubes (4). A vast majority of these studies have emphasized elaboration of mechanisms governing efficient incorporation of nucleotides (speed) with high specificity (fidelity). Whether or not speed is a consequence of fidelity, vice versa, or completely separable is not clear (17, 33, 52-54). However, these two parameters are linked to recombination efficiency. Moreover, the studies reported herein point to the existence of a biochemical property other than speed and fidelity, referred to here as undefined, that contributes to recombination efficiency. Together, the observations reported herein demonstrate that the current state of the art precludes association of a single biochemical property to viral fitness, virulence, and/or pathogenesis.

## Materials and Methods

### Cells and Viruses

Adherent monolayers of HeLa and L929 fibroblasts were grown in DMEM/F-12 media. Media was supplemented with 100 U/mL penicillin, 100 µg/mL streptomycin, and 10% heat inactivated (HI)-FBS. All cells were passaged in the presence of trypsin-EDTA. Wild-type and recombinant PV viruses were recovered after transfection of RNA produced *in vitro* (see below) from full-length cDNA or from the CRE-REP assay parental partners (22). PV type 1 (Mahoney) was used throughout this study (Genbank accession number: V01149.1).

### Plasmids, in vitro transcription, cell transfection, and virus quantification

All mutations were introduced into the pET26Ub-PV 3D (50) bacterial expression plasmids using overlap extension PCR. The presence of the desired mutations and the absence of additional mutations were verified by DNA sequencing. The expression plasmid encoding the various mutations was digested and inserted into an intermediate plasmid, pUC18-BglII-EcoRI-3CD (referred as pUC-3CD in (51), and then the fragment between BglII and ApaI was cloned into the subgenomic replicon, pRLucRA, or viral cDNA, pMovRA or PV1ΔCRE. PV1ΔCRE is a full-length PV type 1 (Mahoney) cDNA bearing 8 synonymous substitutions in the oriI *cis*-acting replication element (CRE) located in the 2C-coding region and was described previously (22). In order to create a fluorescence-based assay for recombination in cells, we modified the PV1ΔCRE acceptor RNA to include the coding sequence for UnaG green fluorescent protein. Briefly, UnaG-encoding sequence (35) carrying a 3C protease cleavage site at its carboxyl terminus was inserted between the IRES and the P1 region of the PV sequence. Translation occurred from the natural poliovirus initiation codon. Proteolytic cleavage and release of the fluorescent protein occurred by normal 3C protease activity. Plasmids encoding PV genomes (full length or subgenomic) were linearized with ApaI. All linearized cDNAs were transcribed *in vitro* using T7 RNA Polymerase and treated with 2U DNAse Turbo (ThermoFisher) to remove residual DNA template. The RNA transcripts were purified using RNeasy Mini Kit (Qiagen) before spectrophotometric quantification. Purified RNA in RNase-free H_2_O was transfected into either HeLa or L929 fibroblasts using TransMessenger (Qiagen). Virus yield was quantified by plaque assay. Briefly, cells and media were harvested at 2-3 days post-transfection, subjected to three freeze-thaw cycles, and clarified. Supernatant was then used on fresh HeLa cells in 6-well plates; virus infection was allowed to continue for 30 min. Media was then removed, and cells were washed twice with PBS (pH 7.4) washes before a 0.8% (w/v) agarose-media overlay was added. Cells were incubated for 2-3 days and then fixed and stained with crystal violet for virus quantification.

### Luciferase assays

Subgenomic luciferase assays were performed as described previously (32).

### Virus sequencing

Viral RNA was extracted from clarified culture supernatant using a Qiagen RNAeasy Mini kit, reverse transcribed using Superscript II reverse transcriptase (Invitrogen) using an oligo-dT primer according to the manufacturer’s protocol. PCR amplification of the 3D^pol^ region in PV used template cDNA and appropriate oligonucleotides as listed in Table 1 by using Phusion high-fidelity DNA polymerase (NEB) according to the manufacturer’s protocol. PCR products were gel purified and sequenced by the Genomics Core Facility of the Pennsylvania State University.

### One-step growth analysis

HeLa cells in 12-well plates were infected by each virus at a MOI of 10. Following a 30-minute incubation, cells were washed twice with PBS and media was replaced. Virus was harvested at different time-points post infection and the virus yield was quantified by plaque assay.

### Quantitative RT-PCR

Viral RNA was purified from virus stocks by using QiaAmp viral RNA purification kit (Qiagen) and used for RT-qPCR to determine genome copies. This analysis was performed by the Genomics Core Facility of the Pennsylvania State University. DNAse-treated RNA was reverse-transcribed using the High Capacity cDNA Reverse Transcription kit (Applied Biosystems, Foster City CA) following the protocol provided with the kit. Quantification by real-time qPCR was done with 2X TaqMan Universal PCR Master Mix (Applied Biosystems, Foster City CA) in a volume of 20 μL, with primers 5′-ACCCCTGGTAGCAATCAATATCTTAC-3′(forward) and 5′-TTCTTTACTTCACCGGGTATGTCA-3′ (reverse), and probe 5′-[6-Fam] TGTGCGCTGCCTGAATTTGATGTGA-3′ in a 7300 Real-Time qPCR System (Foster City CA) machine. A standard curve was generated using in vitro transcribed RNA.

### Ribavirin-sensitivity assay

This assay was described previously (48). HeLa cells were treated with 600 µM ribavirin for 1 h before infection. Ribavirin-treated cells were then infected at MOI 0.1 with each virus variant. Following infection, the cells were washed with PBS and media was replaced with ribavirin. Infection was allowed for 24 h. Cells and supernatant were subjected to three freeze-thaw cycles. Media was clarified and used for plaque assays. All yields were normalized to an untreated control.

### Cell-based recombination assay

This assay was developed in the laboratory of David Evans and was used as described previously (22, 34). L929 fibroblasts were transfected with the PV donor and accepter RNAs, both carrying the same mutations in the RdRp gene. For the fluorescence-based detection of recombinants, the UnaG green fluorescence reporter was encoded by the acceptor RNA. Supernatant at 2∼3 days post-transfection was used to infect HeLa cells. Recombinant viruses were either quantified by plaque assay or by fluorescence imaging (488nm (ex) and 509 nm (em)).

### Polymerase expression and purification

Purified polymerases used for biochemical analysis were prepared as described previously (50, 51).

### Poly(rU) polymerase activity assay

Reactions contained 50 mM HEPES, pH 7.5, 10 mM 2-mercaptoethanol, 5 mM MgCl_2_, 60 μM ZnCl_2_, dT_15_ (2 μM), poly(rA) (100 μM AMP), UTP (500 μM), [α-^32^P]UTP (0.2 μCi/μL), and PV RdRp (0.2 μM). Reactions were initiated by addition of PV RdRp and incubated at 30 °C for 5 min at which time the reactions were quenched by addition of EDTA to a final concentration of 50 mM. Reaction volumes were 50 μL. Products were analyzed by DE81 filter binding, where 10 μL of the quenched reaction was spotted onto DE81 filter paper discs and dried completely. The discs were washed three times for 10 min in 250 mL of 5% dibasic sodium phosphate and rinsed in absolute ethanol. Bound radioactivity was quantitated by liquid scintillation counting in 5 mL of Ecoscint scintillation fluid (National Diagnostics).

### sym/sub-based template switching assay

The sym/sub assay has been described previously (22, 36). Elongation complexes were assembled by incubating 5 μM PV RdRp with 1 μM sym/sub RNA primer-template and 500 μM ATP for 5 min (Mix 1). Template-switching reactions were initiated by addition of 60 μM RNA acceptor template and 500 μM CTP, GTP and UTP (Mix 2) and then quenched at various times by addition of 50 mM EDTA. All reactions were performed at 30 °C in 50 mM HEPES, pH 7.5, 10 mM 2-mercaptoethanol, 60 μM ZnCl_2_, and 5 mM MgCl_2_. Products were analyzed by denaturing polyacrylamide gel electrophoresis, visualized using a PhosphorImager, and the transfer products quantified using ImageQuant TL software (GE Healthcare).

## Acknowledgements

This study was supported by a grant (AI45818) from NIAID, NIH to CEC. AW is the recipient of a fellowship (18POST33960071) from the American Heart Association (AHA).

